# Automation of yeast spot assays using an affordable liquid handling robot

**DOI:** 10.1101/2022.07.16.500326

**Authors:** Shodai Taguchi, Yasuyuki Suda, Kenji Irie, Haruka Ozaki

## Abstract

The spot assay of the budding yeast *Saccharomyces cerevisiae* is an experimental method that is used to evaluate the effect of genotypes, medium conditions, and environmental stresses on cell growth and survival. Automation of the spot assay experiments from preparing a dilution series to spotting to observing spots continuously has been implemented based on large laboratory automation devices and robots, especially for high-throughput functional screening assays. However, there has yet to be an affordable solution for the automated spot assays suited to researchers in average laboratories and with high customizability for end-users. To make reproducible spot assay experiments widely available, we have automated the plate-based yeast spot assay of budding yeast using Opentrons OT-2 (OT-2), an affordable liquid-handling robot, and a flatbed scanner. We prepared a 3D-printed mount for the Petri dish to allow for precise placement of the Petri dish inside the OT-2. To account for the uneven height of the agar plates, which were made by human hands, we devised a method to adjust the z-position of the pipette tips which is based on the weight of each agar plate. During the incubation of the agar plates, a flatbed scanner was used to automatically take images of the agar plates over time, allowing researchers to quantify and compare the cell density within the spots at optimal time points *a posteriori*. Furthermore, the accuracy of the newly developed automated spot assay was verified by performing spot assays with human experimenters and the OT-2 and quantifying the yeast-grown area of the spots. This study will contribute to the introduction of automated spot assays and the automated acquisition of growth processes in conventional laboratories that are not adapted for high-throughput laboratory automation.

## Introduction

The budding yeast *Saccharomyces cerevisiae*, a unicellular eukaryote, is an important model organism in a wide range of research areas, including gene regulatory networks and metabolic pathways [1]. One of the basic experimental procedures in yeast research is a spot assay on agar plates. Spot assays using agar plates are frequently used to assess the effect of genetic mutations on cell growth and the drug resistance of mutant strains [2]. They are also used to detect genetic interactions using synthetic growth defects and protein-protein interactions using the two-hybrid method [3,4]. In a spot assay, researchers spot the yeast culture on agar plates using a micropipette or pin to visually or quantitatively assess the growth capacity and survival based on the density of the cells in spots of the same size [5]. Comparison of the cell densities between strains of different genotypes and under different media conditions allows for biological hypotheses of interest to be generated and tested and for the screening of genetic and environmental factors that influence their growth. However, the experimental procedures of spot assays can be complicated and require skill in preparing a dilution series and spotting on agar media, especially when processing many samples. In addition, the experimenters must check the agar plates frequently during incubation to obtain spot assay results at the time when the differences in the yeast growth become easy to see. These characteristics necessitate multiple instances of human interventions and could result in variation in the results of the spot assays.

By automating the experimental procedures in a yeast spot assay from preparing a dilution series to spotting to observing spots continuously, it is expected to improve the reproducibility and reliability of the results, help allocate human resources to other tasks, and enable high-throughput experiments. The advent of liquid-handling robots has made it possible to automate the preparation of a dilution series and spotting using solid pins, split pins, needles, and tips [6,7]. Similarly, the automation of the observation of colonies and spots on agar plates using continuous imaging with a flatbed scanner has been proposed [8–11] and specialized plate imaging systems [5,7,12]. Indeed, automation of yeast spot assay experiments including spotting and observation has been implemented by combining colony-handling robots or robotic liquid-handling workstations equipped with pin tools, and purpose-built plate photographing systems, especially for high-throughput functional screening assays [13–15]. However, such dedicated automation equipment in laboratory automation is expensive and difficult for the average laboratory to introduce [16]. This hinders many researchers from benefiting from automated spot assay experiments of budding yeast. Moreover, considering that yeast spot assays are used in conjunction with other experiments, the ability to perform spot assays on Petri dish agar plates, which are commonly used in yeast research, has the advantage of being easily coordinated with other experimental procedures that are conducted by humans. Therefore, it is important to design an affordable solution that average laboratories, without a large budget, can easily and reproducibly deploy to make automated experiments widely available to the research community [17–19]. Furthermore, the software and hardware of such automation systems should be easy to customize to adjust to changes in labware and protocols [20,21] and to integrate into larger AI-driven autonomous experimental systems [22–26].

The Opentrons OT-2 (OT-2; Opentrons Labworks Inc., New York, USA) is an automated pipetting machine that is a low-cost and open-source hardware, easy to cutomized by end-users, and is controlled by a populatr programming language Python [27]. The OT-2 is used for automating experimental procedures in yeast research, such as high-throughput screening [28], micro-cultivation experiments [29], DNA cloning [30], bioreactor assays [31], and proteomics [32]. If it is possible to automate yeast spot assays using OT-2, it is expected to be a solution with a low cost and few technical barriers that can be used in various laboratories, especially in academia. However, there are two challenges to automating spot assays using the OT-2. First, because the highly viscous agar medium is decanted onto Petri dish plates, the height of the agar varies among the agar plates depending on the amount of medium that is poured and the degree of drying. While human experimenters are robust to differences in agar height between plates, automated liquid-handling robots require tip height adjustment for each plate during pipetting. In particular, because the yeast suspension solution is more viscous than water, if the tip does not touch the agar medium surface, the water droplets will remain attached to the pipette tip. Thus, to properly spot the yeast suspension solution on the agar medium surface, the tip must be close enough to touch the agar medium surface. The second challenge is the issue of fitting the labware. Labware needs to be fitted in the same position for proper dispensing by the liquid-handling robots. However, unlike Society for Biomolecular Screening (SBS)-format plates that are used in high-throughput experiments [11], Petri dish-type plates cannot be fit into the deck at the bottom of the OT-2, so the Petri dishes need to be adapted to the SBS format [33].

To make reproducible spot assay experiments widely available, we proposed an affordable method for automating yeast spot assays based on Petri dish agar plates using the OT-2 and a flatbed scanner. To deal with the uneven heights of the agar plates, which were made by human hands, we devised a method to adjust the z-position of the pipette based on the weight of each agar plate. In addition, a dedicated mount for the Petri dish was designed and fabricated using a 3D printer to allow for the precise placement of the Petri dish inside the OT-2. To take images of the agar plates over time, a flatbed scanner was used, allowing researchers to quantify and compare the cell density within the spots at optimal time points *a posteriori*. We verified the accuracy of the newly developed automated spot assay by performing spot assays with human experimenters and the OT-2 and quantifying the yeast-grown area of the spots.

## Materials and Methods

### Code availability

The Python scripts, shell scripts, Jupyter notebooks, and 3D CAD data that were used in the experiments and the data analysis are available from the GitHub repository: https://github.com/bioinfo-tsukuba/ot2_spot_assay

### Preparation of YPD liquid medium

Powdered reagents at a final concentration of 2% Peptone (Bacto Peptone, 211677; Thermo Fisher Scientific, Waltham, Massachusetts, US) and 1% Yeast Extract (Bacto Yeast Extract, 212750; Thermo Fisher Scientific) were added to a 500 mL glass bottle and dissolved in 450 mL of deionized water to make up the yeast extract peptone (YP) liquid medium. In parallel, 500 mL of 20% dextrose (D(+)-glucose, 045-31167; FUJIFILM Wako Pure Chemical Corporation, Tokyo, Japan) solution was prepared with deionized water in a 500 mL glass bottle. The opening of each glass bottle was covered with aluminum foil, and it was autoclaved with an autoclave machine (LSX-500; TOMY SEIKO Co. Ltd., Tokyo, Japan) at 120°C for 2 h. After autoclaving, 50 mL of the 20% dextrose solution was added to 450 mL of the YP liquid medium to make up the yeast extract peptone dextrose (YPD) liquid medium.

### Preparation of YPD and YPD+TM agar plates

For the YPD agar medium, powdered reagents at a final concentration of 2% Peptone, 1% Yeast Extract, and 2% Agar (STAR Agar L-grade 01, RSU-AL01-500G; Rikaken Co., Tokyo, Japan) were added to a round flask and dissolved in 450 mL of deionized water. The flask’s opening was covered with aluminum foil, and it was autoclaved with an autoclave machine (LSX-500; TOMY SEIKO Co., Ltd.) at 120°C for 2 h. After autoclaving, 50 mL of 20% dextrose solution was added to 450 mL of the medium to make up the YPD agar medium.

For the YPD and Tunicamycin (YPD+TM) agar medium, 500 mL of the YPD agar medium was prepared in a round flask as described above. The flask was cooled to about 60°C by stirring with a hotplate stirrer (HPS-2002; LMS Co. Ltd., Tokyo, Japan) for 15 min, then, the aluminum foil lid was removed while on a clean bench (MCV-131BNF; PHC Holdings Corporation, Tokyo, Japan). Next, Tunicamycin (202-08241; FUJIFILM Wako Pure Chemical Corporation) was added at a final concentration of 1 μg/mL, and the opening was again covered with aluminum foil and stirred for about 5 min.

Then, the YPD agar or YPD+TM agar medium was poured onto Sterile Veritable Petri Dishes (Shallow, 36-3412; IWAKI (AGC), Shizuoka, Japan) on a clean bench. The Petri dishes were left on a clean bench for 2 d to allow the agar to harden and the surface to dry.

### Yeast strains

Two yeast strains were used in this study: a wild-type (WT) strain (*MAT***a** *ade2 trp1 can1 leu2 his3 ura3*) and a *hac1*Δ mutant strain (*MAT***a** *ade2 trp1 can1 leu2 his3 ura3 hac1*Δ*∷kanMX6*). These yeast strains were generated in a previous study [34].

### Experimental design of the spot assay

To compare the OT-2 and human experimenter spot assays, we designed a comparison experiment using the WT strain and the *hac1*Δ strain as a control, as it is known not to grow at all on the YPD+TM agar medium [35]. Within the same plate, spotting was repeated three times for each strain to validate the variability of the spotting procedure. To confirm the variability between plates, two plates of the same type were prepared for each YPD agar and YPD+TM agar medium. In this experiment, the dilution factor was set to 5x. Therefore, each dilution series consisted of five spots. Seven dilution series were made on each plate: dilution series 1, 2, and 3 were the WT strains; dilution series 4 was spotted with autoclaved deionized water (H_2_O) as a negative control; and dilution series 5, 6, and 7 were the *hac1*Δ strains (**Fig. 3B**).

### Preparation of the yeast strains

The yeast strains from −80°C freeze stock were inoculated on the YPD agar medium and were incubated in a 30°C incubator (MIR-154-PJ; PHC Holdings Corporation) for 2 d.

### Liquid culture

In autoclaved test tubes, 3 mL of the YPD liquid medium was added, and the WT and *hac1*Δ strains were each inoculated. The test tubes were incubated in a shaking water bath (Personal-11, 0069409-000; TAITEC CORPORATION, Saitama, Japan) at 30°C and 200 rpm for 16 h. Then, 300 μL of culture medium in each of the test tubes was inoculated in 3 mL of fresh YPD liquid medium in a new autoclaved test tube, and they were incubated in a shaking water bath at 30°C and 200 rpm for 3 h.

### Spot assay executed by a human experimenter

#### Measuring the absorbance of the yeast cultures

Initially, 180 μL of H_2_O was placed in wells A1, A2, A3, A4, A5, A6, and A7 of a 96 well plate. Next, after 3 h of incubation of the WT and *hac1*Δ culture medium, 20 μL of WT culture medium was added to wells A1, A2, and A3 of the 96 well plate, 20 μL of H_2_O was added to well A4, and 20 μL of *hac1*Δ culture medium was added to wells A5, A6, and A7 with a yellow tip (AG-200-FP-Y; FCR and BIO Co. Ltd., Kobe, Japan). The absorbance was measured at 620 nm with Absorbance 96 Plate Reader (Byonoy GmbH, Hamburg, Germany). Then, to bring the absorbance of the 10-fold diluted culture to 0.020, the dilution ratio was calculated based on this absorbance.

#### Preparation of the dilution series for spotting

A human experimenter mixed the diluent with H_2_O to reach an absorbance of 0.020 at 620 nm in the first well of the dilution series. The second to fifth wells of each 5-fold dilution series were filled with 160 μL of H_2_O. Then, 40 μL was aspirated from the first well, dispensed into the second well, and mixed by pipetting twice in the second well, and this was repeated up to the fifth well.

#### Spotting on the agar plate

As soon as one dilution series was made, 5 μL from the fifth well was aspirated with a 200 μL yellow tip and spotted onto two YPD agar plates and two YPD+TM agar plates. After the spotting was completed in the first row (one sample), a second dilution series was made and repeated until seven rows (seven samples) were completed.

### Spot assay executed by the Opentrons OT-2

#### Estimation of the agar height by the agar weight

To estimate the agar height from the agar weight, the agar weight was substituted into the following formula:

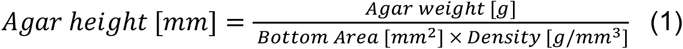

where the diameter of the bottom of the Petri dish is 86 [mm]; the agar density is 0.00107 [g/mm^3^] (yeast extract 5 g, peptone 10 g, dextrose 10 g, and agar 10 g in 500 ml of water); and the weight of the Petri dish (without agar medium) is 17.88 [g]. Therefore, the agar height was calculated as follows:

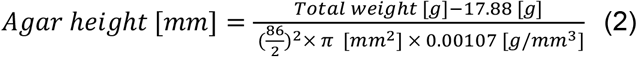

Based on the calculated agar height, we set the tip height of the OT-2 (**Fig. 2A**). When setting the labware calibration of the OT-2, the top of the Petri dish mount that was placed on the deck was set as the bottom of the Petri dish mount (z = 0). Where Z = [the bottom thickness of the Petri dish (2 mm)] + [the agar height calculated by equation (1 mm)] + [the clearance (1 mm)]. The height of the point of a tip was set at z = Z mm to adjust the tip height for each agar medium with a different height. To validate the agar height that was obtained from the calculations, we prepared 16 agar-filled Petri dishes and measured the agar height with calipers at five points—four around the agar and one in the center.

#### Creating labware with a 3D printer

Two types of labware were made using a 3D printer (Finder; Zhejiang Flashforge 3D Technology Co. Ltd., Jinhua, Zhejiang, China). The Computer-Aided Design (CAD) data was uploaded to GitHub (see Code availability). The first jig was a Petri dish mount. We created four Petri dish mounts to mount the Petri dish onto the OT-2 deck. Initially, we measured the outer diameter of a 90 mm Petri dish with a caliper and created a 3D model using Fusion 360 (Autodesk, Mill Valley, CA). Next, we sliced the STL model using Flashprint (Zhejiang Flashforge 3D Technology Co. Ltd.) and printed four of them using a 3D printer. The filament was a FLASHFORGE PLA Filament 0.07 inches (1.75 mm; Zhejiang Flashforge 3D Technology Co. Ltd.). The second jig was a tube rack. We created a tube rack for the OT-2 to handle 15 mL tubes (2324-015N; IWAKI (AGC)) and 50 mL tubes (2324-050N; IWAKI (AGC)). Initially, we used the drawings for the 15 and 50 mL tubes [36,37] as references to create a 3D model using Fusion 360. Then, we sliced the STL model using Flashprint and printed it using a 3D printer. The filament used was a FLASHFORGE PLA Filament 0.07 inches (1.75 mm).

#### Liquid-handling robots

The OT-2 (999-00047, Opentrons Labworks Inc.) was used. The P20 GEN2 single pipette and P300 GEN2 single pipette were set.

#### Setting the labware in the deck of the Opentrons OT-2

We placed labware on the OT-2 deck as follows (**Fig. 1B**): Deck 1, Opentrons 300 μL tips (Opentrons Labworks Inc.); Deck 2, tube rack (a 50 mL tube filled with H_2_O was put into the hole labeled as A1); Deck 3, Corning 96-well polypropylene microplate (3364; Corning Inc. NY, US); Deck 4, a tips rack for Opentrons 20 μL tips (Opentrons Labworks Inc.); Deck 5, YPD agar plate 1 on the Petri dish mount; Deck 6, YPD agar plate 2 on the Petri dish mount; Deck 8, YPD+TM agar plate 1 on the Petri dish mount; and Deck 9, YPD+TM agar plate 2 on the Petri dish mount.

#### Executing the spot assay with the Opentrons OT-2

The automatic spot assay program was executed using a laptop computer (MacBook Air, Apple Inc., CA, US) connected to the OT-2 via Secure Shell (SSH). Specifically, the bash script named “run_spotassay_readABS_yeakiest_biological-test.sh” was executed in the terminal application. When the bash script started, the config.csv was opened and the experimenter entered (1) the number of samples (number of dilution series), (2) the number of agar plates that were being tested, (3) the starting position of the p300 tip rack, (4) the starting position of the p20 tip rack, and (5) the weight of the agar plates with a plate lid (In this research, YPD agar plate 1: 41 g, YPD agar plate 2: 38 g, YPD+TM agar plate 1: 42 g, YPD+TM agar plate 2: 48 g). After entering the data, the human experimenter saved the config.csv and typed yes into the terminal application, then the OT-2 began to run, and dispensed 180uL of H_2_O into the wells of the 96-well plate on Deck 3 according to the number of samples.

#### Measurement of the absorbance of the yeast cultures

After the OT-2 dispensed 180 μL of H_2_O into wells A1, A2, A3, A4, A5, A6, and A7 of the 96 well plate, the human experimenter added 20 μL of the WT strain culture to wells A1, A2, and A3, 20 μL of H_2_O to well A4, and 20 μL of the *hac1*Δ culture to wells A5, A6, and A7. Then, we measured the absorbance at 620 nm using Absorbance 96 Plate Reader via the vendor-provided GUI software and saved the measurement as a CSV file named OD620.csv. Next, when “d” was typed in the terminal, “recipe_spotassay.py” was executed, and the dilution rate was calculated to adjust the absorbance at 0.020 among the samples (A1–A7), with the calculation result saved as “recipe_spotassay.csv”.

#### Preparation of the dilution series and spotting with the Opentrons OT-2

The files recipe_spotassay.csv and spotassay.py were transferred to the OT-2 via SSH, and the OT-2 executed the spot assay protocol. The OT-2 protocol is available on GitHub (see “Code availability”). This OT-2 protocol includes the following experimental procedure: (1) make the first well in a dilution series by dispensing *x* μL of a 10x dilution of the yeast culture and 200-*x* μL of H_2_O, where *x* is defined in recipe_spotassay.csv; (2) dispense 160 μL of H_2_O from the second to the fifth wells in a dilution series; (3) take 40 μL from the *i*th well, dispense to the *i*+1th well, and mix it twice, repeat this for *i* = 1, 2, 3, and 4; (4) take 5 μL of the *j*th well and spot it onto the four agar plates on the decks 5, 6, 8, and 9, repeat this for *j* = 5, 4, 3, 2, and 1; and (5) when one dilution series is completed, repeat the procedure from the beginning to make a second dilution series.

### Time-lapse scanning of the agar plates

To quantitatively evaluate the spot assay results, images of the agar plates are often taken using a digital camera or scanner [5]. Since budding yeast has various growth rates and lag time depending on the genotype and culture conditions [38], human intervention is usually required for spot assay experiments, such as conducting preliminary experiments in advance to determine the time when the pictures of the agar plates need to be taken or checking the plates frequently while culturing. However, recent studies have proposed image acquisition of agar plates through automatic control of a scanner to reduce human intervention in observing the agar plates [8–11]. Therefore, we constructed an automatic system for acquiring images of agar plates over time (**Fig. 1C**) and determined the appropriate time for selecting an image for quantification according to various genotypes and culture conditions.

In this system, a flatbed scanner (GT-X980; Seiko Epson Corp., Japan) was placed in an incubator and scanned at certain time intervals according to a shell script (see “Code availability”). Since it is known that budding yeast divides once every 90 min when incubated at 30°C [39,40], the automated scanning was performed at a time interval of once every 90 min.

After the spot assay was completed by a human experimenter and the OT-2, the agar plates were placed on the scanner in an incubator at 30°C and oriented to scan from the back side of the spot surface. The incubator door was gently closed to prevent the plate from shifting due to vibration. A program was run on a desktop computer (HP EliteDesk 800 G5 SF, HP Japan Inc., Tokyo, Japan; Ubuntu 20.04 LTS) that was connected to the scanner that initiated scanning at a resolution of 600 dpi every 90 min; the program was interrupted by pressing Ctrl+C when the display in the Terminal reached more than 90 scans.

### Quantification of the relative yeast growth

Yeast growth on agar plate can be quantified by area of colonies: The total cell count and the colony area generally show a linear relationship within a certain range of inocula [41]. To quantify the area, we measured the mean gray value of each spot using ImageJ Fiji [5]. For each region of interest (ROI) and all the time points, the mean gray value (which is the average of the gray values of all the pixels in an ROI) was calculated for each spot. In ImageJ, the gray value is calculated by converting each pixel to grayscale using the formula: gray = 0.299*red + 0.587*green + 0.114*blue.

Using Python pandas, the mean gray value was normalized by dividing the mean gray value for each spot at each time by the mean gray value at 0 min. This normalized value was defined as the relative yeast growth.

### Data analysis

Visualization and data analysis of the relative yeast growth data was performed using Python. The OLS function from the statsmodels package was used for fitting the regression line, and the NumPy package was used to calculate the Coefficient of Variation (CV) of the relative yeast growth for each medium, strain, and plate. The plots were drawn using the plotnine package.

## Results and Discussion

### Development of the labware for the automatic yeast spot assays using the Opentrons OT-2 liquid handling robot

We designed two types of labware to automate the yeast spot assay experiments on the OT-2, with one being a Petri dish mount. Many laboratories that handle microorganisms use round Petri dishes to keep the microorganisms, but the decks of the OT-2 are designed to hold SBS-format labware, so the round Petri dishes cannot be placed on the OT-2. To hold a round Petri dish for spotting using the OT-2 pipettes, we designed and created Petri dish mounts according to the SBS standard to be set in the deck at fixed positions (**Fig. 1A**).

**Figure 1.**
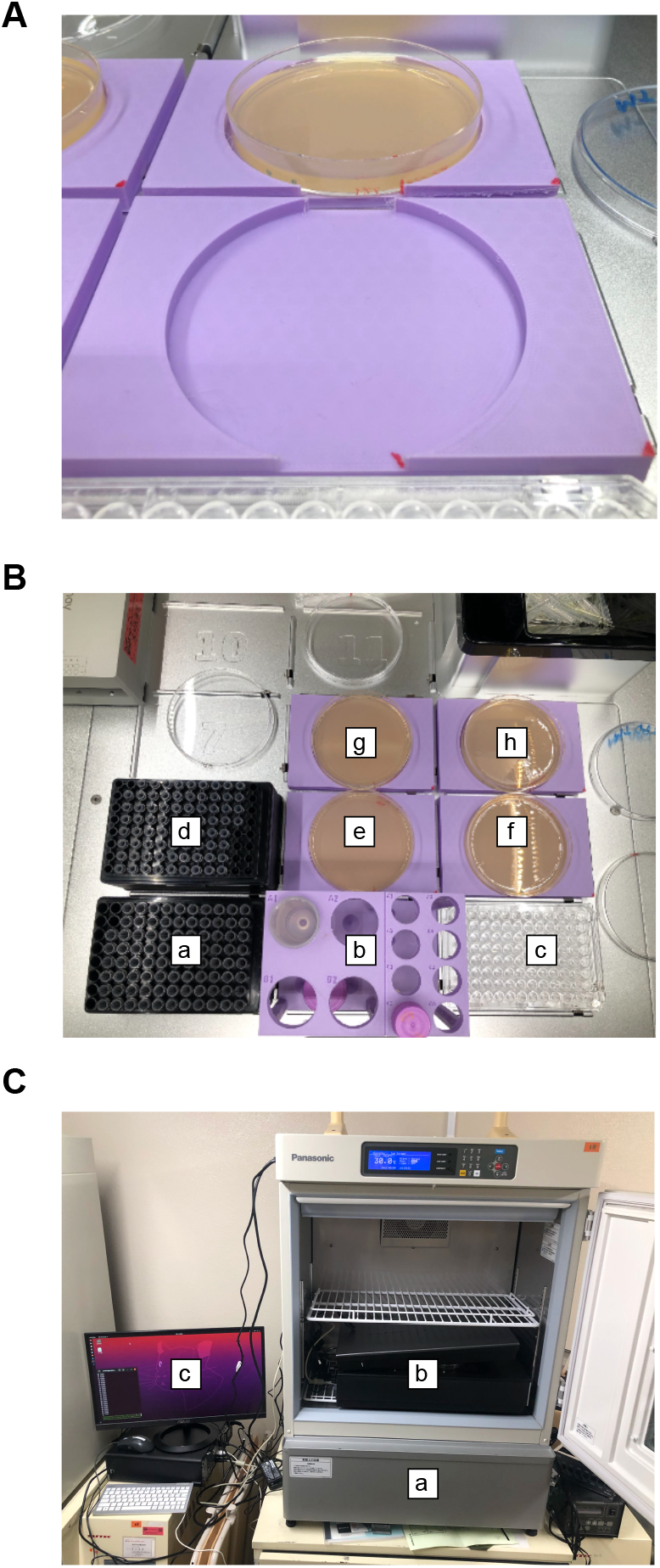
Implementation of an automated yeast spot assay on agar plates. (A) Petri dish mount. (B) Configuration of the Petri dish mounts and other labware inside the Opentrons OT-2. Each small letter indicates: a, p300 tip rack; b, tube rack (50 ml tube is set into location A1); c, 96-well plate; d, p20 tip rack; e, YPD plate on the Petri dish mount; f, YPD plate on the Petri dish mount; g, YPD+Tunicamycin (TM) plate on the Petri dish mount; and h, YPD+TM plate on the Petri dish mount. (C) Scanner set in an incubator. Each small letter indicates: a, incubator; b, scanner; and c, computer.

The second type of labware was a tube rack. For the spot assay experiment that used the OT-2 in this study, approximately 10 mL of H_2_O was consumed in total when measuring the absorbance and preparing the dilution series of the yeast cultures. At the time of development, we used 12 well plates but they needed to be filled with H_2_O every time the H_2_O ran out. To solve this problem, we designed an SBS-format tube rack using a 3D printer so that the OT-2 could handle H_2_O in 50 mL tubes and also modified the protocol to lower the tip by 0.1 mm each time the H_2_O was aspirated, which led to a robust solution for the lack of water.

The two designed labware and other labware were placed on the decks of the OT-2 to perform the automatic spot assay (**Fig. 1B**).

### Adjustment of the tip height according to the agar weight

A successful spotting with micropipettes necessitates contact dispensing [6]: the point of the tip should be close enough to the agar surface so that it almost touches the agar when spotting onto the agar (**Fig. 2A**). When using agar plates made by human experimenters in the automatic spot assay, the problem was that the height of the agar plates varied slightly depending on the amount of the agar that was poured into a Petri dish and the dryness of the agar. When human experimenters perform spotting, they can recognize the height of the agar with their eyes, so they can easily shift the point of a tip closer to the appropriate height and spot the agar. However, since the OT-2 does not have a sensor to detect the height of the agar plates, it is difficult for an OT-2 with the default configuration to robustly accommodate agar plates with different heights. To solve this problem, we devised a method to adjust the height of the point of a tip according to the agar height by calculating the height based on the agar weight.

**Figure 2.**
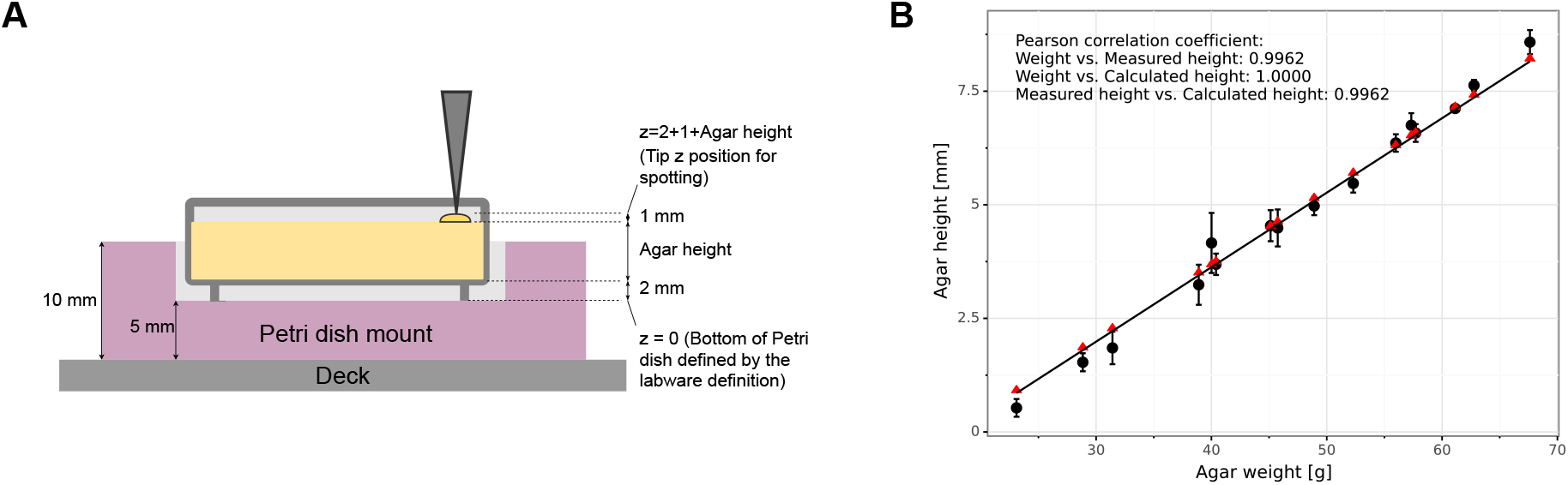
Estimation of the surface height based on the weights of the agar plates. (A) Schematic of the parameters of the plate heights. A Petri dish is located on a Petri dish mount on the Opentrons OT-2 deck. (B) Relationship between the agar weight [g] and agar height [mm]. Each black circle and error bar represents the mean and standard deviation of the five measurements of agar height for each plate. Each red triangle represents the calculated agar height of each plate. The Pearson correlation coefficients between the weights, measured heights, and calculated heights are shown.

To verify the agar height that was estimated using Equation (1), the actual agar height was measured and compared with the estimated value (**Fig. 2B**, **Supplementary Table 1**). The Pearson correlation coefficient between the measured and estimated agar height was 0.9985 (**Fig. 2B**), confirming the validity of the method for calculating the agar height from the agar weight. The tip height of the OT-2 was adjusted to correspond to the estimated agar height. This allowed us to bring the point of a tip close enough to touch the agar (**Fig. 2A**).

### Evaluation of the spot assay plates between a human experimenter and the Opentrons OT-2

To evaluate the developed automatic spot assay system with the OT-2, we conducted a spot assay executed by a human experimenter and the OT-2 and compared them qualitatively and quantitatively (**Fig. 3A**). We designed the following spot assay experiments: two YPD plates and two YPD+TM plates. The experimental design was performed for the spot assay with humans and the OT-2 (**Fig. 3B**).

**Figure 3.**
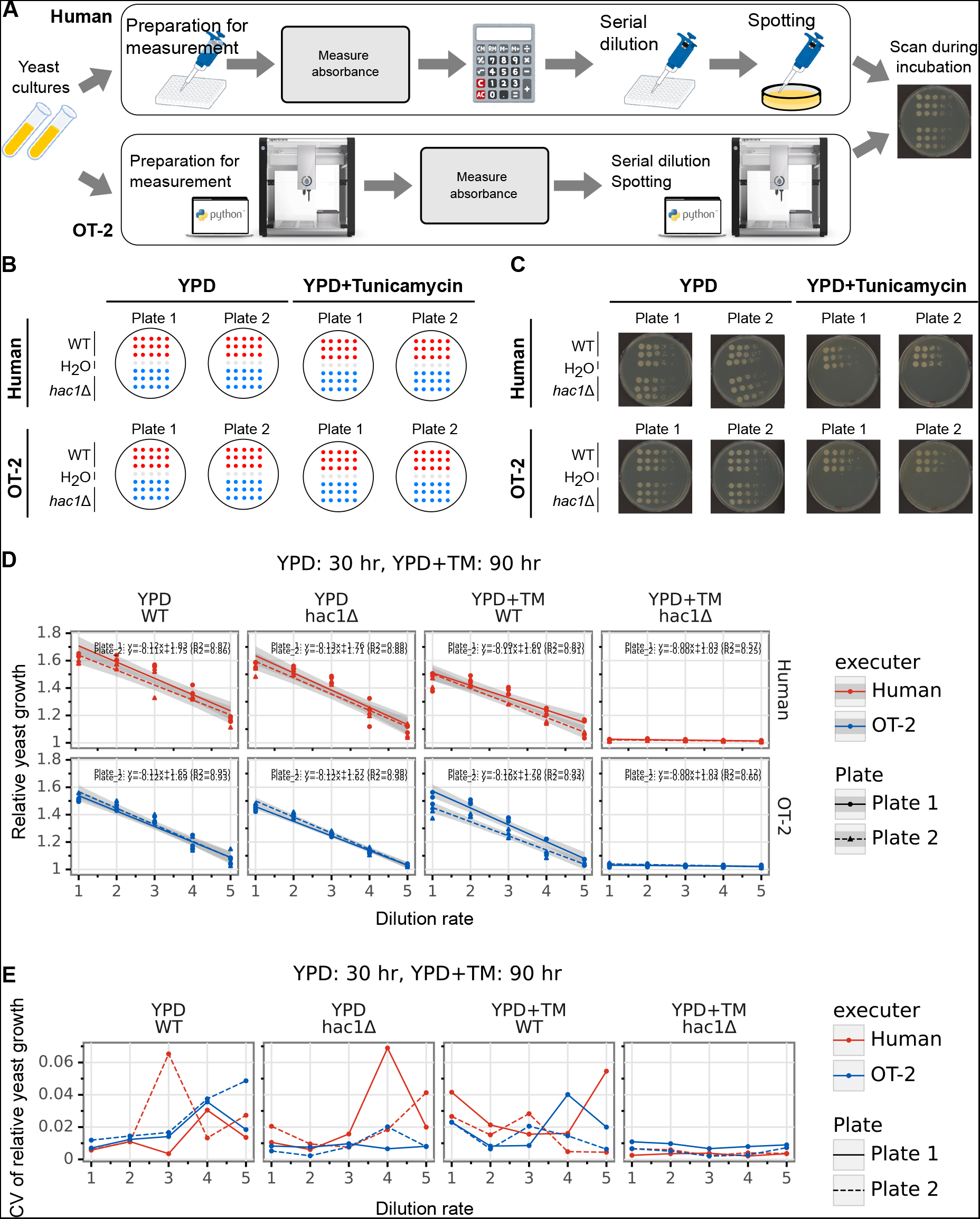
Comparison of the spot assay results conducted by a human experimenter and the Opentrons OT-2. (A) Overview of the spot assay experiments which were performed by humans and the Opentrons OT-2 (OT-2). The spot assay by a human experimenter consisted of (1) preparation of yeast cultures in a 96-well plate for measuring the absorbance of yeast cultures, (2) measurement of absorbance, (3) calculation of dilution factor, (4) serial dilution, and (5) spotting. The spot assay by OT-2 consisted of (1) preparation of yeast cultures in a 96-well plate for measuring the absorbance of yeast cultures, (2) measurement of absorbance, (3) serial dilution, and (4) spotting. The spotted plates were placed on a flatbed scanner in an incubator and scanned during incubation. The OT-2 image was adopted from https://opentrons.com/ot-app/. (B) Schematic of the experimental design. On each agar plate, seven dilution series, each consisting of five spots, were generated: three series were for the wild-type WT strain, one series was for H_2_O, and three series were for the *hac1*Δ mutant strain. Two types of agar plates were used: YPD and YPD+Tunicamycin (TM). Two replicates were prepared for each condition. (C) Scanned images of the agar plates after 30 h (YPD) and 90 h (YPD+TM). (D) Quantification of the area of the spots that were generated by the human experimenter (red) and the OT-2 (blue) after 30 h (YPD) and 90 h (YPD+TM). The x-axis represents the dilution rate (minus the logarithm with base 5). The y-axis represents the normalized area. Solid (plate 1) and dashed (plate 2) lines represent the linear regression lines with the gray area representing confidence intervals. (E) The coefficient of variation (CV) of the normalized area of the spots that were generated by the human experimenter (red) and OT-2 (blue) after 30 h (YPD) and 90 h (YPD+TM). The x-axis represents the dilution rate (minus the logarithm with base 5). The CV was calculated for plates 1 (solid lines) and 2 (dashed lines) separately.

The *HAC1* gene is a key regulator in the endoplasmic reticulum stress response pathway in budding yeast, and it is known that the *hac1*Δ mutant strain exhibits a significant growth defect on YPD+TM plates [35]. Therefore, it was expected that we would observe no difference in the relative growth between the WT and *hac1*Δ mutant on the YPD plates, but we would observe smaller relative growth of the *hac1*Δ mutant than that of the WT on the YPD+TM plates.

To evaluate the cell growth on the agar plates, the plate images were taken using a flatbed scanner (**Fig. 1C**), and the images were compared at 30 h for the YPD plates and 90 h for the YPD+TM plates. Visual inspection between the human experimenter and OT-2 plates showed similar growth between the three replicates, consistent with the expected phenotype in both the human experimenter and OT-2 plates. The cell growth of the WT and the *hac1*Δ mutant on the YPD plates were similar between the three replicates, plates, and human experimenter and OT-2, and yeast growth decreased with the dilution rate. However, on the YPD+TM plates, the WT showed growth that was comparable to that on the YPD plate, while the *hac1*Δ mutant did not grow at all (**Fig. 3C**), which was consistent with previous studies [35].

Next, for the quantitative evaluation, the relative yeast growth was quantified using the scanned images of the agar plates and we conducted linear regression with the dilution rate of each plate (**Fig. 3D**, **Supplementary Table 2**). The slopes of the linear regression were less than 0 for the WT on the YPD and YPD+TM plates and the *hac1*Δ mutant on the YPD plates for both the human experimenter and OT-2. In addition, all the plots of the *hac1*Δ mutant on the YPD+TM plates had a slope of 0 and no colonies grew. These results are consistent with the expected phenotype. To further confirm the reproducibility among the three spot series for each condition, we calculated the CV of the relative yeast growth for each different spot at the same dilution rate. There was no significant difference in the CV between the OT-2 and human experimenter (Mann–Whitney U test, *p*<0.05 after the Bonferroni correction) (**Fig. 3E**, **Supplementary Table 3**). These results indicate that the yeast growth on the agar plates was comparable between the human experimenter and OT-2.

### Potential limitations and future directions

In this study, we developed an automated yeast spot assay using the OT-2 and confirmed that no differences were observed between the experimental results of the human experimenter and the automated system. Our proposed method can perform the entire process including the dilution series preparation, spotting, continuous observation of yeast growth, contributing to improved reproducibility of experimental manipulations, reduction of human errors, and traceability of results. However, this automated system has several potential limitations that should be mentioned. The first is the throughput of the experiment. The automated spot assay that was developed in this study can spot seven different samples per plate, and only a maximum of six plates can be placed onto the decks of the OT-2 per experiment. Thus, for the current configuration, the number of spot assays that can be performed at one time is limited to a maximum of 42 series, which is lower than the use of manual pin tool [5]. To improve the sample-size throughput, it is possible to increase the number of samples per plate by (1) reducing the liquid volume of spotting and increasing the density of spots and/or (2) using an SBS-format 96-well and 384-well plate. The second is the time that is required for the experiment. The system developed in this study took about 25 min to spot four plates, while the experimental time when it was conducted by the human experimenter was about 30 min. To reduce the time that is required for the experiment, it would be best to optimize the movements [42] and dispensing speed of the OT-2 pipette and replace the single pipette with multichannel pipettes. The third is the quantification of the relative yeast growth. The mean gray value was used in this study, however, the number and size of the colonies are often used to measure the growth potential of yeast. Several methods have been proposed to quantify the number and size of colonies from scanned images [8,9], and the use of these methods could also be a possible solution. The fourth is the statistical power of the spot assay experiment. Since we used the *hac1*Δ mutant strain, which shows similar growth to the WT on YPD plates and no growth on YPD+TM plates, further investigation is needed to verify how much difference in growth potential can be detected. Detailed analyses of the statistical power of the time-lapse images of agar plates would provide the opportunity to quantify the difference in the growth potential that is hard to distinguish by qualitative visual inspection [5,43] and to clarify the effects of genetic mutations and culture conditions on the proliferation in more detail [8,44].

The affordable yeast spot assay automation system that was proposed in this study can be used, not only in yeast genetic and stress response research but also in synthetic biology [45], which is expected to develop substantially in the future. By using the present method, reproducible spot assay results can be easily obtained at optimal timing, even in small laboratories [16]. Moreover, since the OT-2 hardware and software are open source and can be controlled by Python, a popular programming language for AI development, our poposed system is expected to be easily integrated into future AI-drivin autonomous experiment system [22–26]. This research is expected to contribute to the promotion of decentralized laboratory automation by lowering the barrier to entry for laboratory automation in extensive wet labs.

## Acknowledgments

We thank Mamoru Suzuki, Takaaki Horinouchi, Ai Muto, and Tomoaki Mizuno for their technical advice and thoughtful discussions. We would like to thank Editage (www.editage.com) for English language editing.

## Supplemental Material

**Supplementary Table 1 Original data for Fig. 2B**.

**Supplementary Table 2 Original data for Fig. 3D**.

**Supplementary Table 3 Original data for Fig. 3E**.

